# Use of machine learning for quantification of retinal pigment epithelium tight junctions improves assay sensitivity

**DOI:** 10.1101/2024.05.24.595580

**Authors:** Yan Gao, Mark-Anthony Bray, Michael Twarog, YongYao Xu, Natasha Buchanan, Yiyun Zhang, Quintus Medley, Magali Saint-Geniez, Ganesh Prasanna, Qin Zhang

**Affiliations:** Ophthalmology Research, Novartis BioMedical Research, Cambridge, Massachusetts, USA; Discovery Sciences, Novartis BioMedical Research, Cambridge, Massachusetts, USA

**Keywords:** Retinal pigment epithelium, Tight junction, Confocal microscopy, Automated quantification, Image analysis, open-source software, Machine learning

## Abstract

The retinal pigment epithelium (RPE) is critical for maintaining outer retinal barrier homeostasis. In age-related macular degeneration (AMD), the RPE can undergo a dedifferentiation process that includes tight junction (TJ) loss and displacement of zonula occludens-1 (ZO-1), which may impair structural and functional integrity of the RPE barrier and contribute to disease pathogenesis. Our objective was to develop an automated and sensitive quantification method for TJ aberrations in an RPE immunofluorescence imaging assay, following treatment with TNFα or TGFβ2. However, quantifying ZO-1 morphological changes in the RPE using standard image analysis methods did not provide a satisfactory assay window. To address this challenge, we developed an imaging assay to quantify ZO-1 changes using a machine learning approach, enabling enhanced phenotypic characterization of the ZO-1 changes in RPE cells and improved assay sensitivity. We were also able to capture and quantify the reversal of these changes using etanercept, an TNFα inhibitor, with this imaging assay. Our findings indicated that this machine learning ZO-1 quantification assay could serve as a potential phenotypic readout for RPE dedifferentiation and enabling large-scale mechanistic studies.

## 1. Introduction

Retinal pigmented epithelium (RPE) is responsible for maintaining the functional integrity of both photoreceptors and the choroidal vasculature (Lakkaraju et al., 2020). Loss of RPE differentiation is known to play a critical role in numerous retinal diseases, including AMD (Ban and Rizzolo, 2000; Shu et al., 2020; Zhou et al., 2020). Dedifferentiation can also lead to epithelial-mesenchymal transition (EMT), a transdifferentiation process consisting of complex cellular and molecular changes that can result in loss of cell junctions and aberrant RPE migration (Sripathi et al., 2021; Zhou et al., 2020). The RPE is composed of a single layer of cells joined laterally towards their apices by tight junctions (TJ) between adjacent plasma membranes. TJs mediate size-selective passive diffusion of solutes to and from the outer segments of the retina (Naylor et al., 2019). Junctional integrity of human-induced pluripotent stem cell-derived RPE (iPS-RPE) cells can be visualized by staining the scaffold protein zonula occludens-1 (ZO-1). Under normal conditions, matured RPE cells exhibit a characteristic cobblestone morphology with intact cell borders delineated by continuous TJs. Induction of EMT in iPS-RPE following treatment with TNFα or TGFβ2 results in disruption of the TJ structure, leading to the translocation of TJ-associated proteins such as ZO-1. Hence, quantification of the TJ structural changes can provide insights into overall RPE health and functionality (e.g., low permeability) and may serve as a marker for detecting dysfunction in more complex mechanisms related to transport, metabolism, or polarity.

Measurement of the transepithelial resistance is the canonical approach to measure RPE barrier function *in vitro*, but has limited throughput, is time-consuming, and may overlook spatially-resolved information about regional differences in barrier properties within a tissue (Napoli and Strettoi, 2023). In contrast, image-based quantification of RPE cells using microscopy holds promise for conducting high-throughput studies (Ye et al., 2020). Imaging of ZO-1 phenotypes may provide an orthogonal approach for assessing RPE health, especially for studies where large numbers of perturbations are under consideration, permitting observation of single-cell changes in TJ morphology to complement barrier function assessment.

That said, the alteration of the cell boundaries and distribution of morphological changes in RPE-progressive diseases like AMD are highly heterogeneous, with individual RPE cells exhibiting a variety of cytoskeletal phenotypes. Such heterogeneity in RPE morphological changes lays in both the variety of responses triggered by various stressors, and by the multiple phenotypes spatially distributed across a sample that a single stressor may produce (Tarau et al., 2019). To address this challenge, we combined deep-learning cell identification with a machine learning approach to identify individual disrupted cells versus unperturbed phenotypic subpopulations, thereby improving assay sensitivity and enabling mechanistic RPE studies at a larger scale.

## 2. Materials and Supplies

### 2.1 Cell culture

iPS-RPE (#R1102; Fuji Film Cellular dynamics) were plated in 384-well plates (#781092; Greiner Bio-One) at a seeding density of 250K cells/cm^2^ and cultured in RtEGM media (#00195409; Lonza Bioscience) with 2x/weekly media changes for 10 days. On day 10, the media was changed to XVIVO-10 media (#BP04-743Q; Lonza Bioscience) and cells were maintained in culture with 2x/weekly media changes. Cells matured for at least three weeks were used to perform experiments.

### 2.2 Cell treatment

To obtain a TNFα dose response curve, RPE cells were grown as specified above and were treated with TNFα (#210-TA; R&D Systems), at 1:2 serial titrations with 12 concentrations starting from 400 ng/mL for 24 hours before cell fixation. In some experimental conditions, cells were treated with 1 ng/ml TGFβ2 (#302-B2; R&D Systems) for 8 consecutive days with replenishment of TGFβ2 every other day then the cells were fixed at day 8. For the dose dependent inhibition experiment, the cells were treated with 200 ng/mL TNFα, immediately followed by exposure to the TNFα blocker etanercept (ENBREL, Amgen) at 1:3 serial titrations starting from 10 µg/ml, 12 points, four biological replicates, and incubated for 24 hrs. Cells were then fixed and stained as described below.

### 2.3 Immunohistochemical Staining

Cells were fixed with 4% paraformaldehyde (#15714-S; Electron Microscopy Science) for 20 min at room temperature (RT) and washed three times with phosphate buffered solution (PBS). The fixed cells were blocked with 5% goat serum with 0.1% Triton X-100 in PBS for 1 hr, stained with mouse anti-human ZO-1 (#3391100; ThermoFisher) at 5 ng/ml at 4°C for overnight, and Alexa Fluor488-goat anti-mouse (#A21121; Life technology) at 1:500 dilution for 45 min at RT. ZO-1 images were acquired using an ImageXpress confocal microscope (Molecular Devices) at 4 non-overlapping fields of view at 40x magnification; a maximum intensity projection was created from 22 Z-slices acquired with a 0.6 µm spacing.

### 2.4 Barrier function assessment

RPE cells were initially seeded at 200K cells/cm^2^ in transmembrane wells (#353095; Corning) and grown as previously specified. The 24-well plate density of RPE post maturation was approximately 1800K cells/cm^2^ or 600K cells/well. Barrier function was assessed by monitoring transepithelial resistance (TER) every 15 min by means of a cellZscope 2 (NanoAnalytics GmbH, Münster, Germany). Baseline TER values (100-200 Ω/cm2) and polarization were comparable to those reported for adult human RPE cultures and native tissue (Blenkinsop et al., 2015). Resistance values for individual monolayers at specific times were normalized to baseline resistance prior to stimulation (as 100%).

### 2.5 Statistical Analyses

All statistical analyses were performed with TIBCO Spotfire Analyst 12.0.7 (TIBCO Software). The Z’ (Z-prime) factor, a common metric of assay quality, was calculated as the statistically robust version of the usual formula (Zhang et al., 1999):

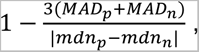

where the median absolute deviation (*MAD*) and the median (*mdn*) of the positive (*p*) and negative (*n*) controls are used instead of the standard deviation and mean, respectively. The Z’-factor is as a measure of the separation between the positive and negative controls: values between 0.5 and 1 indicates an excellent assay, values between 0 and 0.5 indicate a possibly acceptable assay, and values less than 0 indicate the assay is of low quality.

## 3. Detailed Methods

### 3.1 Cellular and tight junction (TJ) quantification

Examples of acquired ZO-1 images are shown in Figure 1 for TNFα and TGFβ2-treated and untreated cells. Post-acquisition, the ZO-1 images were quantified with CellProfiler open-source cellular image analysis software (Stirling et al., 2021). The workflow was as follows: the CellProfiler workflow protocol (“pipeline”) was loaded, and the acquired images served as input to the pipeline. Automated processing of each ZO-1 image, identified single cells and their associated ZO-1-positive TJs. Well-level aggregate values were generated by calculating the median of the single cell measures and served as the output for downstream analysis.

**Figure 1:**
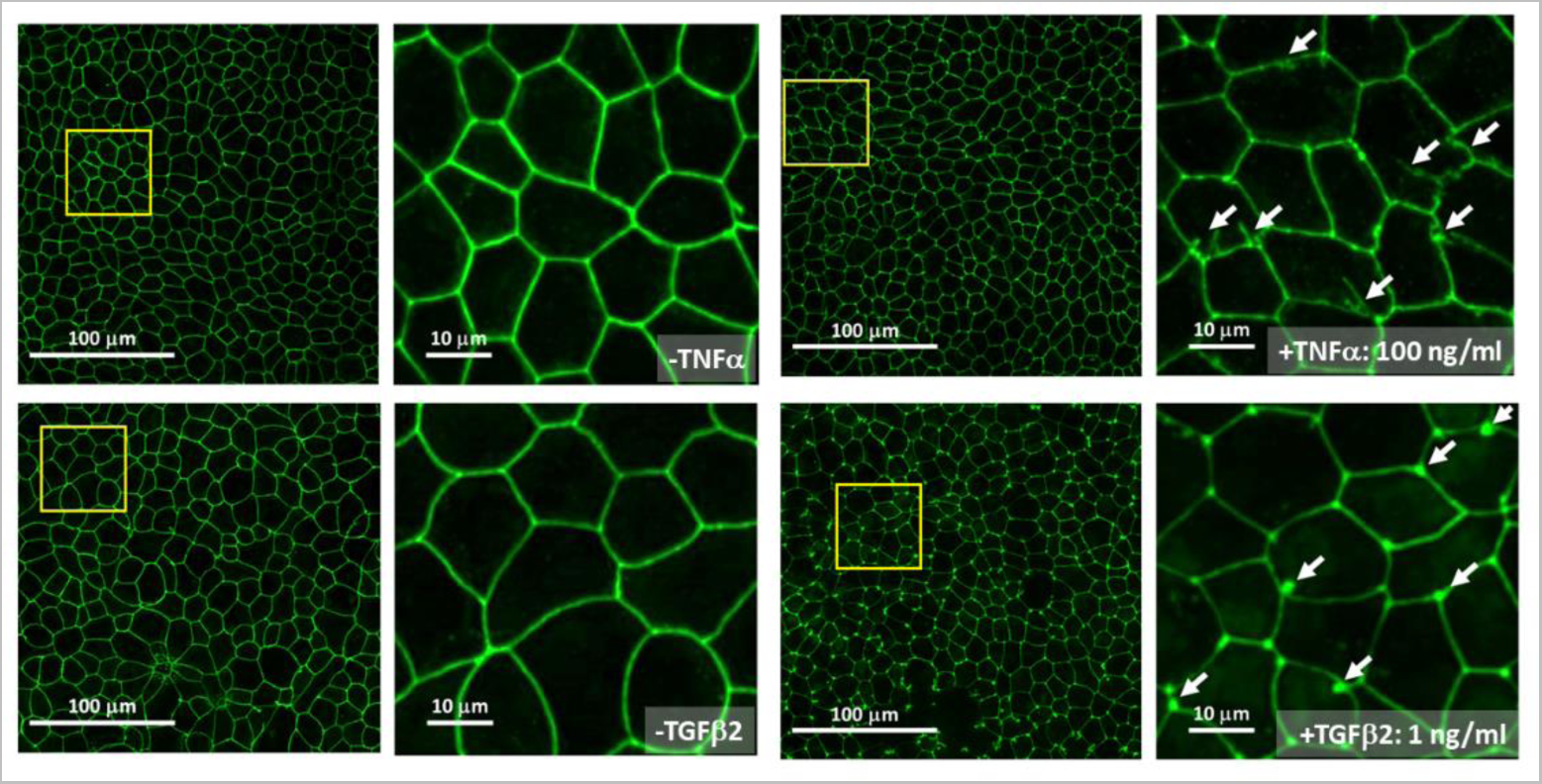
Immunofluorescence microscopy images of ZO-1 staining in RPE cultures. Examples of TNFα- and TGFβ2-induced morphological changes are shown. White arrows indicate examples of disrupted tight junction (TJ) morphology, with the increasingly disrupted “zig-zag” TJs of TNFα-treated cells at and punctate TJs for TGFβ2-treated cells.

For cell segmentation (i.e., thresholding for positive cell staining, followed by delineating the boundaries between touching cells), the watershed approach included with CellProfiler is often successful for typical cellular assays. Unfortunately, this approach to cell identification was not ideal: the cell boundaries were unable to be consistently delineated due to heterogeneity in the ZO-1 staining (Figure 2A). To address this issue, we applied the Cellpose plug-in, a deep-learning based cellular segmentation tool (Stringer et al., 2021), available at https://github.com/CellProfiler/CellProfiler-plugins. Implementing this approach yielded better TJ segmentation (Figure 2B), but interestingly also decreased the Z’ factor for several typical morphological features (Table 1).

**Figure 2:**
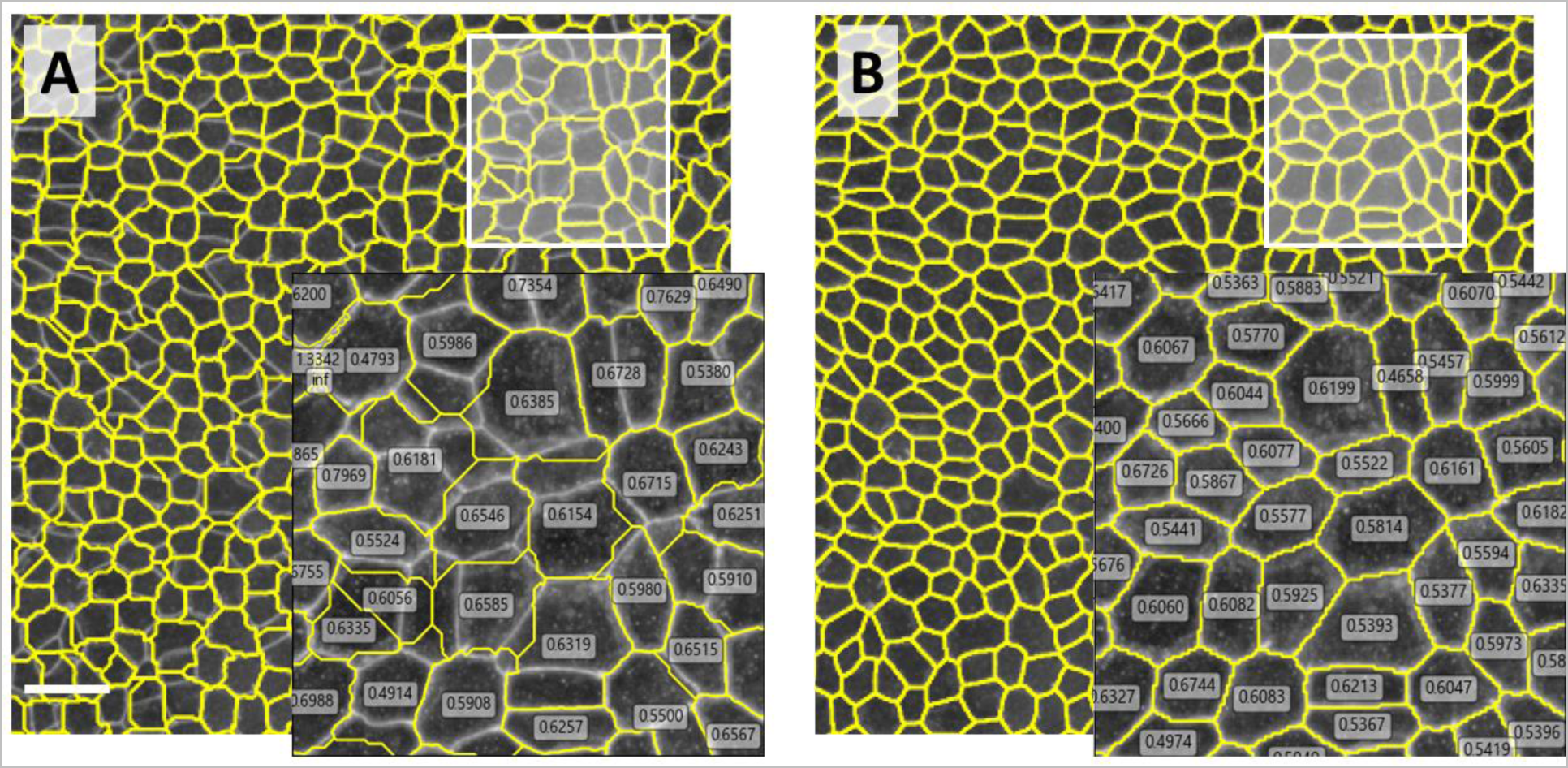
Inaccuracies in cell border identification impact downstream cellular readouts. (A) Using the standard CellProfiler cell segmentation watershed approach (shown as yellow outlines) leads to incorrect calculation of cell morphology, such as the form factor (numeric labels shown in inset). (B) Using the Cellpose approach yields improved segmentation and associated form factor readouts. Scale bar: 40μm.

**Table 1:** Comparison of Z’ factors for several representative CellProfiler features measuring cell morphology, the zig-zag index, and the classified readouts from supervised machine learning. Eccentricity: ratio of the distance between the foci of the ellipse and its major axis length; values range between 0 (circle) and 1 (line). Form factor: 4π area/perimeter^2^; equal to 1 for a circle. Solidity: [object area] / [convex hull area]; Median radius: mean distance of any pixel in the object to the closest pixel outside of the object. Dashes: not calculated.

| Feature name | Watershed | Cellpose |
| --- | --- | --- |
| Area | -0.678 | -2.674 |
| Perimeter | -1.358 | -13.81 |
| Eccentricity | -0.370 | -0.833 |
| Form factor | -0.077 | -0.247 |
| Solidity | -0.005 | 0.419 |
| Median Radius | 0.679 | -1.134 |
| Zig-zag Index | -- | -0.074 |
| Supervised classification (cells) | -- | 0.753 |
| Supervised classification (TJs) | -- | 0.808 |

Initial assay quantification was performed using the “zig-zag index” (Tokuda et al., 2014). This approach requires the researcher to create a sampling of both straight and disrupted TJs manually; the zig-zag index is then defined as the ratio of the freehand lines (*L_Fh_*) to that of straight lines (*L_St_*), i.e., *L_Fh_*/*L_St_*. Since the large scale of the study precluded manual annotation, we used the Cellpose-enabled CellProfiler workflow to recapitulate the ZZI automatically, along with other morphological features. However, the improved cell segmentation (enhanced TJ detection) resulted in a low quality assay (-0.0742; see Table 1), indicating that even with improved cell segmentation (enhanced TJ detection), this metric could not be used for a high-quality assay.

We hypothesized that the low assay quality was due to the presence of a heterogeneous ZO-1 phenotype caused by the compound treatment, i.e., the cell population for a specific treatment consisted of a mixture of straight and disrupted TJs (Figure 3A). These differences would be diluted when taking the median to calculate a well-level aggregate value, resulting in a decreased separation between controls. To improve the assay window, we proposed defining the phenotypes at the single-cell level using a machine-learning approach to construct a classifier.

**Figure 3:**
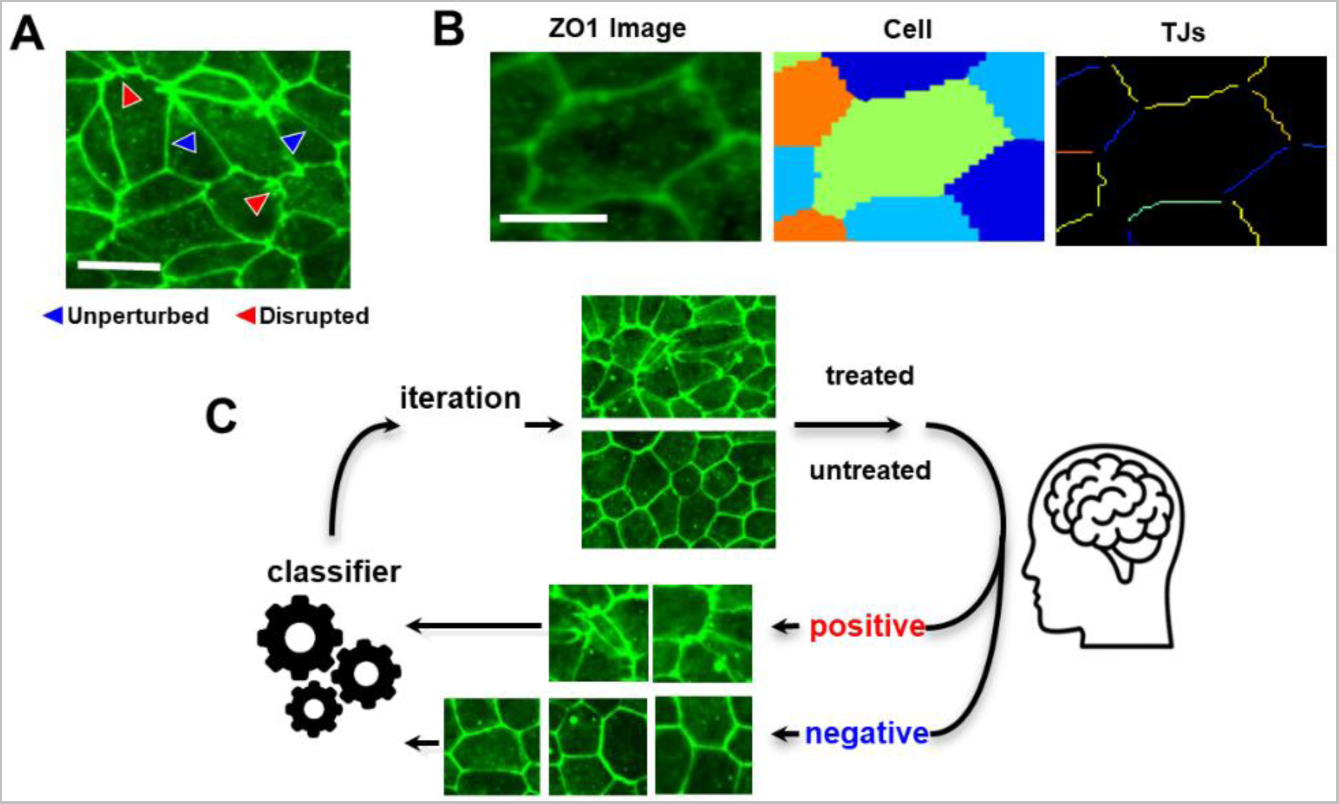
Illustration of the Image analysis and supervised machine learning workflows. (A) An example ZO-1 image highlighting the heterogeneity of TJ phenotype, with both unperturbed and disrupted TJs, Scale bar: 20μm. (B) A CellProfiler/Cellpose-based pipeline is used to accurately identify the ZO-1-delineated cells and TJs, as well as collect single-cell and single-TJ readouts. Scale bar: 10μm. (C) Create a classifier based on the single-cell/single-junction readouts to distinguish the disrupted (”positive”, TNFα treated) junctions or cells from the untreated (”negative”) junctions or cells.

### 3.2 Supervised cellular and TJ classification

The image analysis protocol was revised to generate the appropriate input for a classifier; the new workflow resembled the original one but differed in two ways. First, using the improved cell segmentation, we identified not just the cells but also the TJs themselves (Figure 3B). Second, rather than measuring a handful of morphological features, we instead measured a variety of image-based features (e.g., shape descriptors (Bhatia et al., 2016; Emde et al., 2022), intensity, texture, adjacency statistics) to form a morphological signature. These features were collected for both the cells and TJs. Supervised machine learning was then used to perform classification of the ZO-1 phenotypes (Figure 3C). In this approach, the researcher was tasked with training a predictive model using labeled data to accurately classify new unseen data based on the patterns observed in the training data. Briefly, an initial “training set” was created by the researcher by randomly sub-sampling a collection of objects (i.e., cells or TJs) cultured with the maximum treatment concentration and exhibiting a disrupted phenotype; these objects were assigned a “positive” label for their respective TNFα or TGFβ2 treatment. Similarly, a corresponding collection was sampled from untreated objects (i.e., exhibiting an unperturbed phenotype) and assigned a “negative” label. The researcher was presented with the respective samples in the form of image tiles displaying each object in a graphical user interface. Since the treatment was not uniformly penetrant for all objects in each well, the researcher refined the training set by re-assigning negative-phenotype / positive-labeled object tiles to the negative label using the user interface, and vice-versa for those positive-phenotype / negative-labeled cells. Following a round of label re-assignment, the tool then took the training set of labeled objects (and their associated signatures) and built a classifier capable of distinguishing the positive examples from their negative counterparts. The researcher could then request that the classifier retrieve unsampled objects with the positive and/or negative phenotype and continued to iteratively perform the training procedure to further refine the classifier accuracy. The accuracy of the classifier was evaluated with each iteration with an area under the ROC curve metric (AUC) indicating the probability of the trained classifier correctly distinguishing a positive object from the negative class, or vice-versa. This value ranges from 0 and 1, where 1 indicates perfect prediction, 0.5 indicates random chance, and 0 means the classifier made completely opposite predictions. With this phenotype, the researcher was able to achieve an AUC score of 0.7 or higher (considered to be of acceptable accuracy or better (Hosmer,Jr. et al., 2013)) within 10 iterations. Once sufficient accuracy had been achieved, the researcher then used the trained classifier to label all the objects as positive or negative. The final readout was the normalized number of objects per well classified as positive, i.e., the number of positive-classified objects per well divided by the total number of objects per well. Using the classification results as the new readout yielded a Z’-factor of 0.75 for the cells and 0.81 for the TJs (see Table 1), indicating excellent assay quality.

To confirm that this approach successfully distinguishes the disrupted phenotypes from different treatments, cells were treated with and without TNFα and TGFβ2, and separate classifiers were built for the positive phenotype in each treatment. Figure 4 shows the normalized count (defined as the number of cells of a given class divided by the total cell count) for the two treatments; a similar performance is achieved in both cases, with ∼70% of the cells expressing the TNFα-positive phenotype for the TNFα-experiment and a similar proportion for the TGFβ2-positive phenotype in the TGFβ2-experiment. These morphological differences in TJ staining may be representative of divergent molecular pathways, as EMT induction by TNFα and TGFβ2 has been reported to involve different metabolic responses in RPE (Ng et al., 2023). Corresponding TER data similarly indicates that the treatments may result in functional differences, with TNFα (Supplemental Figure 1A) showing a more substantial decrease in barrier function compared to TGFβ2 (Supplemental Figure 1B). These findings highlight the ability of the ZO-1 machine learning metric to quantify structural changes across the selected treatments, providing additional context to alterations that may result in broad RPE barrier dysfunction.

**Figure 4:**
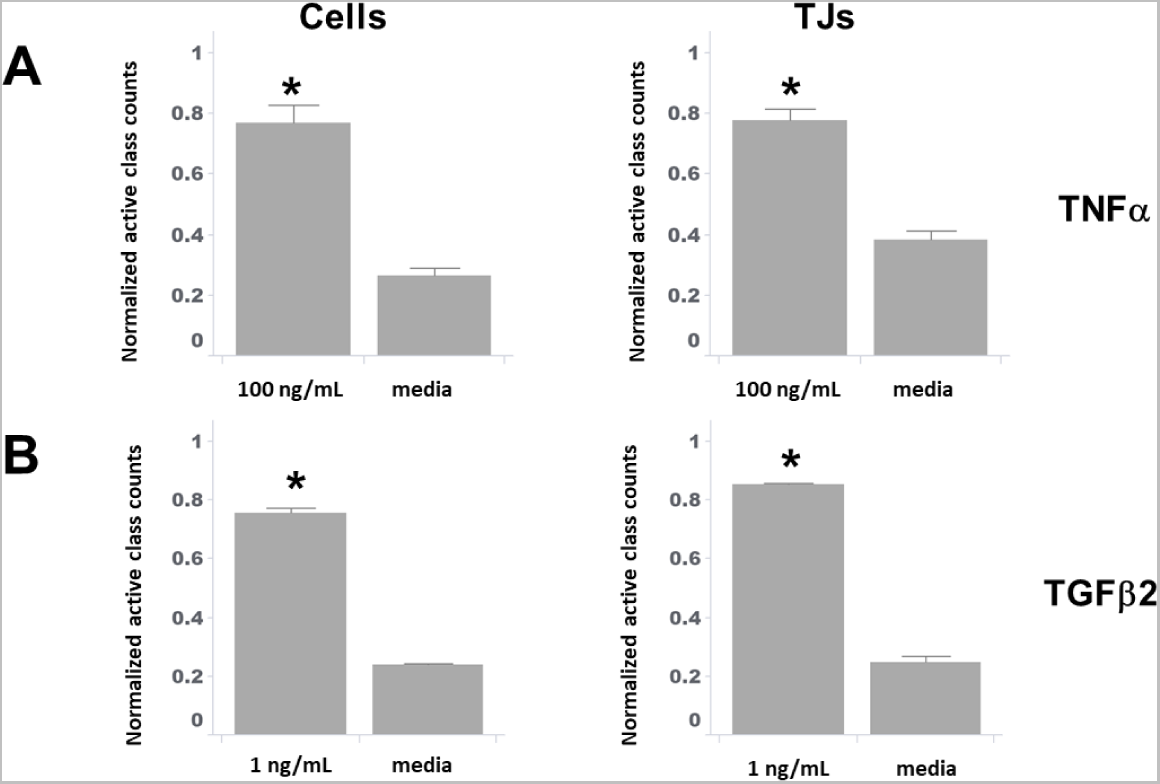
Quantification of a ZO-1-feature classifier to capture morphological changes in iPS-RPE. Classifiers were trained to distinguish the disrupted phenotype prevalent in the treated wells from those observed in the untreated (media) wells. A separate classifier was trained for the TNFα and TGFβ2 experiments. Similar classification performance (defined as normalized counts) is achieved for TNFα (A) and TGFβ2 (B) treatment. Morphological changes are captured whether the cells (left column) or tight junctions (TJs, right column) are classified. Normalized count: number of cells of a given class divided by total cell count. *, *p* < 0.05 compared to relative vehicle control groups. Error bars represent standard error of the mean.

We then performed a TNFα dose-response experiment to examine the sensitivity of our machine learning approach, i.e., whether a classifier can capture the gradual change in phenotypic heterogeneity when the treatment is titrated. For reference, we also characterized the RPE barrier integrity using transepithelial resistance (TER) under TNFα titration (see Supplemental Figure 1B for the full time-course of TNFα TER data). Dose-response curves for the functional and phenotypic response as a function of TNFα concentration at 24 hours is shown in Figure 5, with an IC_50_ of 0.13 ng/mL for TER and EC_50_ of 1.23 ng/mL and 1.09 ng/mL for the cell and TJ classifiers, respectively. These results indicate that the functional TER and phenotypic ZO-1 both capture RPE barrier function defects induced by TNFα, which aligns with previous observations (Twarog et al., 2023). Additionally, the machine learning approach allows for reliable quantification of the phenotypic response.

**Figure 5:**
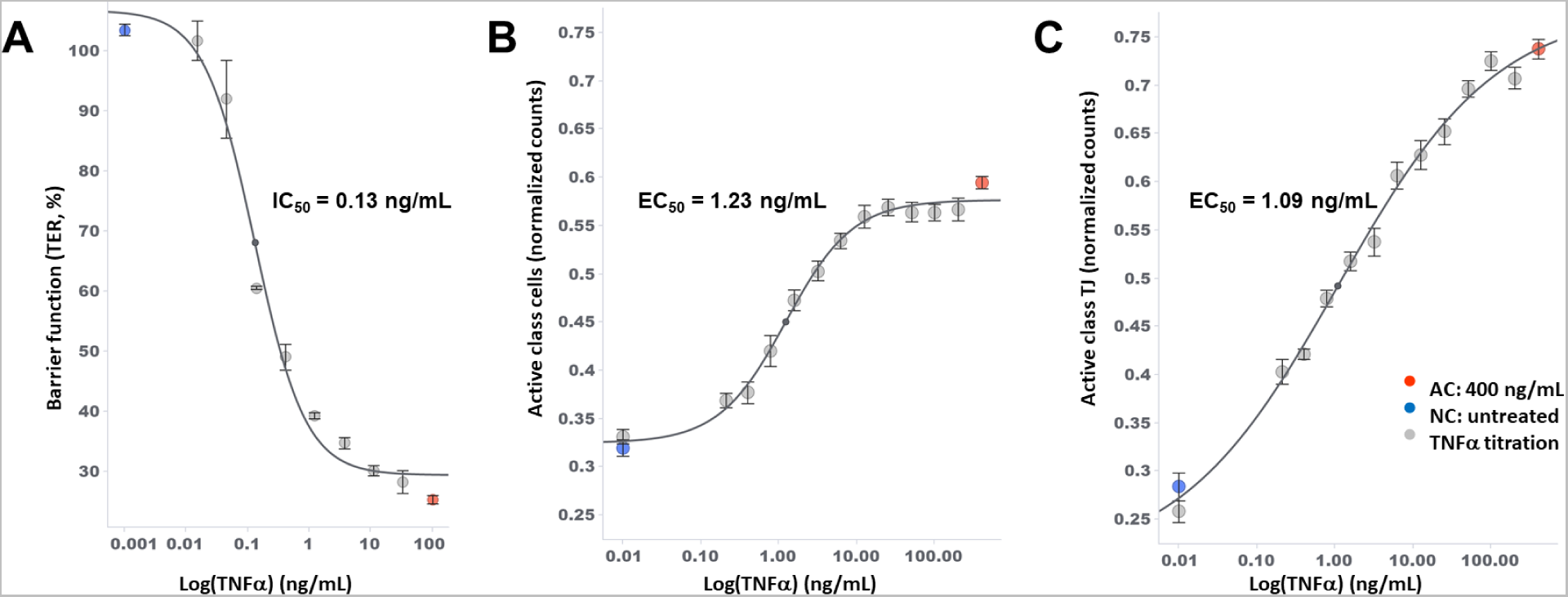
TER and ZO-1 morphological dose-dependent changes post-TNFα treatment in iPS-RPE. Quantification of RPE barrier function as a function of TNFα is shown as TER dose response curve (DRC) (A). In comparison, the DRC of normalized counts highlights the ability of the trained classifier to capture the TNFα-induced changes in ZO-1 phenotype whether the cells (B) or tight junctions (TJs) (C) are classified. Untreated wells or those with the lowest TNFα dose express few cells/TJs classified with the TNFα-positive phenotype, whereas the highest concentration maximizes the number of cells/TJs expressing a TNFα-positive phenotype. Normalized count: number of cells of a given class divided by total cell count. Error bars represent standard error of the mean.

In this study, we utilized a random forest approach for object classification. This method is an ensemble learning technique that combines features to build multiple decision trees for making predictions (Liaw and Wiener, 2002). One advantage of this approach is its ability to calculate the importance of each feature in constructing the classifier. To determine which features were most valuable in building the classifier, we ranked them based on importance. We selected the top seven features for the positive-class cells and TJs, aggregated them to the well level using the median, and calculated the robust Z’-factor (see Supplemental Table 1). Notably, these high-importance features exhibited a low robust Z’-factor at the well level. This observation confirms our initial hypotheses: when used together to construct a classifier, these features can provide a high-quality assay metric at the single-cell level, even if the individual features themselves result in lower assay quality at the well level due to their heterogeneity.

We further validated the assay by examining whether compound treatment could reverse the disorganized phenotype. The TNF inhibitor Etanercept prevents TNFα-induced ZO-1 disorganization in RPE (Goffe and Cather, 2003), and therefore should decrease the proportion of TNFα-positive classified cells/TJs with increasing treatment (Figure 6). For 200 ng/mL of TNFα treatment, the expected dose response was observed, with IC_50_ values derived from the TNFα-positive cells/TJs all within a similar range regardless of classifier.

**Figure 6:**
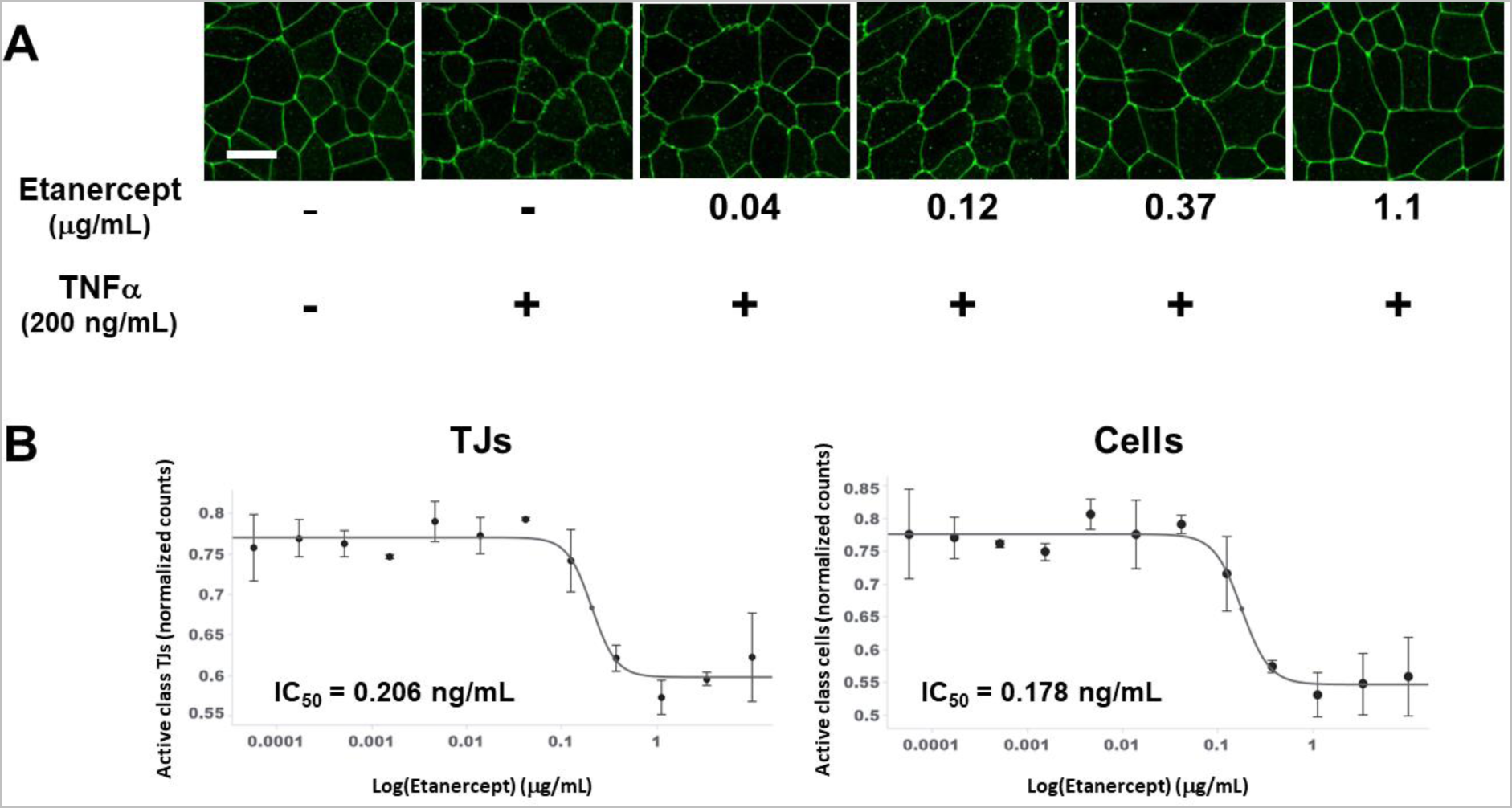
Etanercept prevents ZO-1 disorganization in RPE induced by TNFα at 200ng/mL. (A) TNFα inhibitor etanercept was selected as a tool reagent to treat the cells with a fixed TNFα concentration (200 ng/mL) to obtain the IC_50_ of etanercept. Scale bar: 20 μm. (B) Dose response curves are shown for the TNFα-positive TJ (left) and cell (right) normalized counts, for Etanercept treatments at 200 ng/mL TNFα. Error bars represent standard error of the mean.

## 4. Potential Pitfalls and Troubleshooting

The focus of this paper is on a machine learning quantification method specifically designed to measure the zig-zag phenotype observed in iPS-RPE cells during RPE disorganization, as indicated by ZO-1 TJ staining. However, it is important to acknowledge that there is significant heterogeneity in cytoskeletal phenotypes exhibited by RPE cells in AMD. These include thinning of boundaries between adjacent RPE cells, thickened and frayed RPE borders, enlarged RPE cells, and the presence of multiple intracellular stress fibers (Tarau et al., 2019). Furthermore, these diverse phenotypic features are observed regionally, where different RPE cells from the same donor eye display multiple features in different areas of the RPE. This complexity adds to the challenge of accurately assessing overall cytoskeletal changes in the RPE. Therefore, it is crucial to incorporate the measurement of all these RPE features into the machine learning pipeline to achieve a more precise quantification of overall cytoskeletal changes in the RPE. In addition to the morphological changes in in vivo RPE, TJ changes are also observed when utilizing *ex vivo* ocular flat mount preparations (Boatright et al., 2015; Obert et al., 2017). The deep-learning based segmentation approach is also expected to be applicable to these flat mount samples, provided that the ZO-1 staining is discernable. Additionally, if the changes to the TJs are visually apparent, the supervised machine learning approach should be successful. Evaluation of ocular flat mount samples needs to be performed to verify the theoretical applicability of this ZO-1 machine learning method for quantification of RPE cytoskeletal changes under different and relevant perturbations.

Although we studied the disrupted phenotype induced by TNFα in more detail by using dose-response treatment, the fact that TGFβ2 showed similar results in classifying disrupted vs. unperturbed cells/TJs provides evidence that this approach can be applied to multiple phenotypes, rather than being limited to just one. Even though other phenotypes of ZO-1 disruption can be produced with other treatments (Tarau et al., 2019), we predict that the machine learning approach to phenotypic discrimination remains valid, and the supervised method is expected to also be successful for other treatments, provided the phenotype is visually discernable. In fact, this approach is not limited to only two phenotypes within the same sample. It can also be extended to quantify the enrichment of multiple phenotypes simultaneously (Jones et al., 2009). Additionally, this approach is not confined to RPE cells alone: the quantification of tight junctions can be applied to a diverse range of non-pigmented cells involved in barrier function that express ZO-1, including endothelial cells, epithelial cells, renal tubular cells, among others.

When utilizing a supervised machine learning approach, there are several important considerations to keep in mind. The workflow requires a person to visually assess the phenotype and make corrections to the training set. However, it is important to note that this approach introduces a subjective bias. To ensure its effectiveness, the researcher must have domain knowledge and expertise to make informed decisions about the phenotype. Fortunately, in this case, the positive and negative phenotypes were visually distinct enough that even a non-expert researcher could create an accurate classifier. However, there may be instances where cells appear visually ambiguous, and the researcher is faced with choosing between inappropriate phenotypes for those cells. In such cases, the recommendation is to exclude these ambiguous cells from the training set and focus only on including clear examples of the phenotypes. Ultimately, all cells, including difficult-to-distinguish ones, will be assigned a label. However, either such ambiguous examples are rare and will not significantly affect the results, or a label probability is assigned to account for uncertainty, allowing for the removal of uncertain cells.

The worst-case scenario is that the phenotypes are not sufficiently penetrant to yield a useful result; this situation can be identified during the training step, if the classifier does not return clean examples of the phenotype. Such instances may point back to problems with the biological quality of the assay itself. On the other hand, if the phenotype seems to be visually clear yet the classifier fails to return accurate examples of the phenotype, this may indicate that the collected measurements are insufficient to capture the phenotype. For this reason, it is recommended to collect a wide range of image-based cellular features, and not just those that are hand-selected by the researcher; this approach has been validated for use in unbiased cellular profiling in which the phenotypes are unknown *a priori,* e.g., the so-called “cell painting” assay (Bray et al., 2016). Furthermore, if two stressors, such as TNFα and TGFβ2, produce comparable morphological changes in ZO-1, it becomes necessary to incorporate a second phenotypic assay, such as TER, to better understand the specific impacts on the biological questions and hypotheses under examination. In essence, the automated ZO-1 assay can function independently, or it may need to be combined with another assay to make more informed determinations about target-relevant biology or efficacy endpoints.

To perform the supervised machine learning workflow, our research team used an in-house multi-parametric analysis platform, designed for large-scale usage and scalability. While this enterprise platform is not available for public use, supervised machine learning for single-cell scoring has been previously validated (Jones et al., 2009) and used in the context of ophthalmological research (Dordea et al., 2016). Moreover, the same workflow can be recapitulated with publicly available open-source supervised machine learning tools such as CellProfiler Analyst (Dao et al., 2016; Kornhuber and Dunst, 2022), Enhanced CellClassifier (Misselwitz et al., 2010) and others; a comparison of such tools has been published previously (Smith et al., 2018). While we used the AUC as a classifier accuracy metric on our in-house platform, similar metrics are typically used with publicly available classification tools.

In summary, we have demonstrated a machine learning pipeline that captures the phenotypic changes associated with TNFα induction of ZO-1 morphological alterations in iPS-RPE and rescue of these changes by the TNFα inhibitor etanercept. This machine-learning classification approach enables the automated and unbiased quantification of ZO-1 morphology, suggesting that this ZO-1 imaging assay is a potential phenotypic readout for RPE dedifferentiation and evaluation of overall RPE health. The success of this approach lays the foundation for future phenotypic large-scale studies for the treatment of AMD.

## CrediT authorship contribution statement

**Yan Gao:** conceptualization, investigation, validation, resources, writing – original draft, review & editing. **Mark-Anthony Bray:** investigation, data curation, formal analysis, visualization, writing – original draft, review & editing. **Michael Twarog:** investigation, validation, resources, writing – review & editing. **YongYao Xu:** conceptualization, resources, writing – review & editing. **Natasha Buchanan:** conceptualization, writing – review & editing. **Yiyun Zhang:** investigation, resources, writing – review & editing. **Quintus Medley:** conceptualization, resources, supervision. **Magali Saint-Geniez:** writing – review & editing, supervision. **Ganesh Prasanna:** writing – review & editing, supervision. **Qin Zhang:** conceptualization, writing – review & editing.

## Acknowledgements

The authors thank Pauline Sarraf for discussion of experimental details, and Megan Serpa for ImageXpress confocal microscope maintenance. The authors also thank Novartis BioMedical Research for supporting the research and development.

