## Supplemental material for "Use of machine learning for quantification of retinal pigment epithelium tight junctions improves assay sensitivity"

#### Software installation

The CellProfiler pipeline included with the Supplemental Material was created with version 4.2.4 of the software. To use the Cellpose plug-in (called RunCellpose), it is **necessary to download and install the software from the Python source code** rather than installing the executable available from the download page. This is because the Cellpose plug-in requires additional Python packages not included with the usual executable.

CellProfiler must be installed first before installing the Cellpose plugin-in.

- Documentation on installing CellProfiler from source can be found here for Windows, MacOS/OS X and Linux.
- Documentation on using plug-ins with CellProfiler can be found here, with additional notes about installing RunCellpose here.

If you encounter difficulties, we recommend looking through and/or posting questions on the CellProfiler support forum at Image.sc.

### Transepithelial electrical resistance (TER) measurement

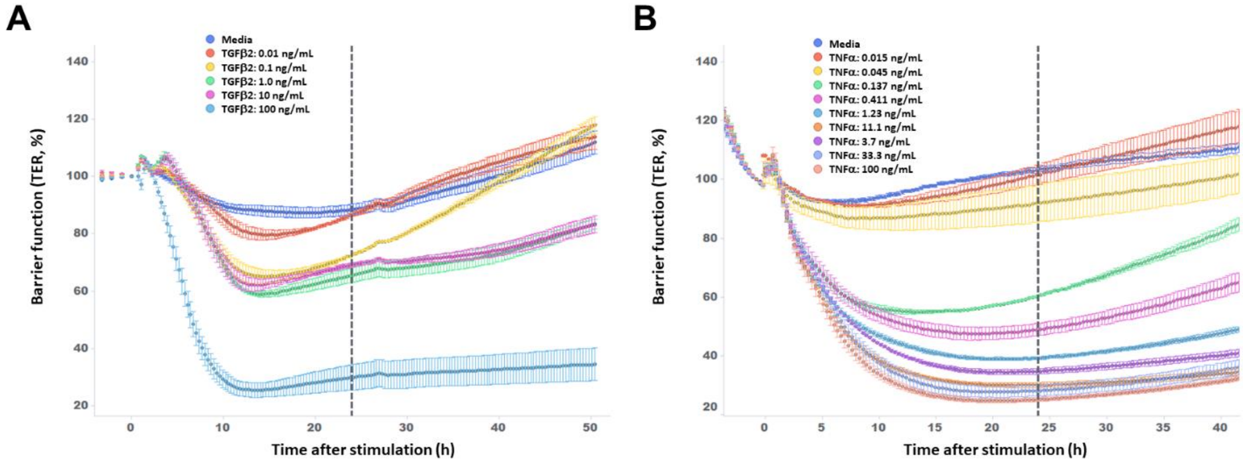


**Supplemental Figure 1: TGFβ- and TNFα-induced impedance loss in iPS-RPE cells.** RPE barrier function was assessed by transepithelial resistance (TER) measurement in response to TGFβ (A) and TNFα (B) titration. Each data point represents biological replicates (n = 3), where error bars represent standard error of the mean. Vertical line: t = 24 hours, the time at which ZO-1 data was collected.

#### Variable importance

| TJ feature name | Robust Z' |
|---|---|
| Intensity_StdIntensity_Distance.ACClass.median | 0.2469 |
| Texture_Correlation_OrigZO1_2_00_256.ACClass.median | 0.2375 |
| Texture_Correlation_OrigZO1_2_02_256.ACClass.median | 0.0837 |
| Intensity_MinIntensity_EnhZO1.ACClass.median | -0.0917 |
| RadialDistribution_FracAtD_OrigZO1_2of2.ACClass.median | -0.1494 |
| Texture_Correlation_OrigZO1_2_01_256.ACClass.median | -0.24 |
| RadialDistribution_RadialCV_OrigZO1_1of2.ACClass.median | -0.3109 |

| Cell feature name | Robust Z' |
|---|---|
| AreaShape_Solidity.ACClass.median | 0.229 |
| AreaShape_Zernike_7_5.ACClass.median | -0.1406 |
| AreaShape_Compactness.ACClass.median | -0.2805 |
| RadialDistribution_FracAtD_OrigZO1_1of4.ACClass.median | -0.3009 |
| RadialDistribution_FracAtD_OrigZO1_2of4.ACClass.median | -0.3584 |
| Texture_Correlation_OrigZO1_5_03_256.ACClass.median | -0.4529 |
| AreaShape_Zernike_8_8.ACClass.median | -0.7174 |

**Supplemental Table 1: Top-ranked features used in the random forest classifier**. The tables list the top 7 image features used in assembling the random forest classifier to distinguish the AC class for the cells (top) and tight junctions (TJ; bottom). These single-object features were aggregated to the well level by taking the median, and the robust Z'-factor was calculated.
